# Optimal antimicrobial response to a changing microbial background at a mucus interface

**DOI:** 10.1101/2023.08.02.551591

**Authors:** Guilherme Volpe Bossa, Shai Bel, Andrew Mugler, Amir Erez

**Affiliations:** Racah Institute of Physics, Hebrew University of Jerusalem, Israel; Institute of Physics and Mathematical Sciences, Southern Chile University, Chile; Azrieli Faculty of Medicine, Bar-Ilan University, Israel; Department of Physics and Astronomy, University of Pittsburgh, USA

**Keywords:** AMP, Host-Microbiome, Muc2, Ileum, Diffusion, SFB, Circadian, Gut

## Abstract

Complex lifeforms host microbiota, microbes that live synergistically with their host. Accordingly, hosts have mechanisms to defend against and tolerate the microbiota. The intestinal mucus, where these systems collide, plays a pivotal role in managing this relationship, yet lacks an integrative theoretical framework. We propose a minimal model to elucidate dynamics at this interface, focusing on the ileum’s mucus defense. The model considers the effect of delay in host antimicrobial peptide secretion and how the host can use two different signals, from the bulk microbiota and from segmented filamentous bacteria (SFB), assuming that the SFB anticipate the bulk microbiota. We propose a theory whereby the host can optimize defense by minimizing antimicrobial peptide production and controlling bacterial exposure. Integrating two recent experiments, we show host dynamics are consistent with sensing both bulk and SFB, supporting our ‘optimal defense’ hypothesis. Therefore, we propose that similar mechanisms could prove advantageous to other species and applicable beyond the ileum’s mucus barrier.

---

A host interacts strongly with its microbiota along the epithelial boundary lining our gastro-intestinal tract: mouth, stomach, and intestines. These boundaries are coated with mucus gels of varying properties, allowing small molecules to move across the mucus while excluding bacteria [1–4]. This mucus is where two worlds collide: on one side of a divide the host epithelium and immune system sit poised, working in harmony and well resourced, but tasked with protecting an enormous boundary—the human gut if stretched would cover a tennis court. On the other side of the divide, a rich community of microbes, the *microbiota*, is engaged in a fierce internal contest for limited resources [5–9]. These two systems sit in close proximity, separated by a mucus layer. Despite the considerable experimental focus on the mucus layer, a comprehensive theoretical framework consolidating these findings remains yet to be established.

Beyond the mucus, in the intestinal lumen, the environment is rich in microbes. On the other side of the divide, the epithelium tolerates only very few or very specific microbes without becoming inflamed [10]. If the lumen microbiota invaded the mucus en masse, coming in contact with the epithelium, an inflammatory response would ensue which could develop to chronic diseases such as IBD [11]. This essential role of the mucus barrier in multiple global health challenges has driven intense research [4, 12–22]. In this study, we aim to clarify a fundamental protective mechanism provided by mucus.

While the protective mechanism of the mucus barrier in the colon is well established, the defense mechanism in the small intestine, is less clear [23]. In the colon, organized as a dense, regular and continuous layer, the mucin *Muc2* forms a firm mesh that physically excludes bacteria from penetrating the dense barrier [2, 3]. However, in the small intestine, though *Muc2* is still abundant, it is not organized in a dense layer as it is in the colon. Indeed, not much is known of the spatial distribution of microbes in the small intestinal mucus layer. Yet, the lumen microbiota are still physically separated from the epithelium cells [24], depicted in Figure 1A. What mechanisms maintain the separation between the epithelium and the microbiota in the small intestine?

**FIG. 1.**
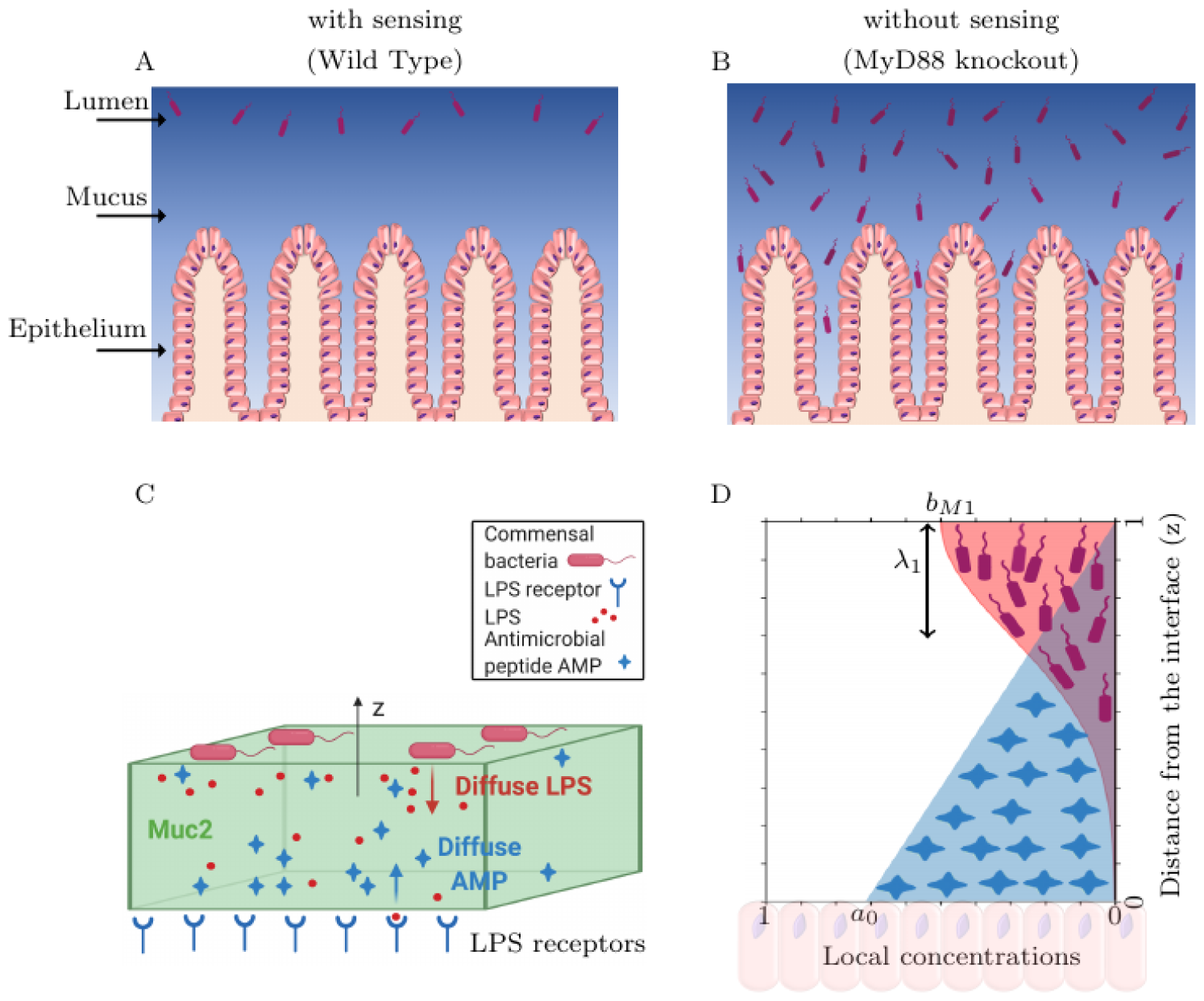
Conjugate-diffusion model of the mucus defense system. Illustration of microbiota localization relative to the ileum epithelium, based on Ref. [24]. (A) In wild-type mice, there is clear separation between the epithelium and the lumen microbiota (red). It was shown that this separation crucially depends on sensing the bacterial signals. (B) Without sensing the bacteria (*MyD88* knockout mice), AMP production is suppressed, the mucus no longer acts as a barrier and the microbiota come in contact with the epithelial cells. (C) Schematic of the conjugate-diffusion model that maintains the ileum homeostasis: bacteria shed cues (e.g. LPS) that epithelial cells detect; upon detection, antimicrobial peptide (AMP) production increases; the AMP diffuse through the mucus and inhibit the bacteria. *z* is perpendicular to the epithelium-microbiota interface, pointing into the lumem, with *z* = 0 the location of the epithelium. (D) Model solution for the local concentrations of a bacterial species (red) and AMP (blue) as function of *z*; Results are using Equation 2 with *b*_*M* 1_ = 1 and the penetration length of the bacteria, *λ*_1_ = 0.3.

Although we propose a general framework for considering host-microbiota interactions through diffusible molecules in the mucus, we have a specific biological system in mind, for which some details now follow. Experiments show that the host defense depends on factors secreted by the enterocytes, paneth and goblet cells [25–28]. Upon sensing the proximity of bacteria, host intestinal epithelial cells release diffusible antimicrobial peptides (AMP). Conversely, when host sensing is inhibited in *MyD88* knockout mice, the bacteria invade [24]. Homeostasis at the host-microbiota intestinal interface changes as the host undergoes diurnal cycles. The cyclic behavior involves bi-directional communication between the host and the microbiota: this communication plays pivotal roles in regulating host immunity and metabolism. For example, recent evidence [29–33] indicate significant variation in host susceptibility to microbial pathogens across the day-night cycle. In recent years it has become evident that the regulation of gut immunity depends on the response induced by bacteria adherent to the small intestine [30, 32, 34]. These adherent, *segmented filamentous bacteria* (SFB), are unique in many ways. The SFB are the only known bacteria to translocate to the epithelium and attach tightly to the epithelial cells in a way that is not pathogenic yet immunomodulatory [35]. Their immunomodulatory effect can be harnessed to make perturbative experiments and can be used to explore physiology during healthy homeostasis [30]. If the SFB anticipate the lumen microbiota, by following the shifts in these adherent SFB bacteria the host may anticipate changes in the microbiota background and synchronize host defenses. Host diet plays a key role [32, 36], as diet changes can undermine the host defenses. For example, subjected to a ‘western-style’ high-fat high-sugar diet, and lacking their preferred nutrients, some commensal species begin to consume the mucus [12, 18]. Furthermore, a high-fat diet disrupts host defense rhythms [32]. Despite significant recent progress in the experimental analysis of the intestinal interface, a deeper understanding of this system is hindered by a lack of theory to integrate the experimental findings [37–39].

What guiding principles keep the microbes in check, preventing them from invading the host? How does the host provide sufficient defense yet avoid excess inflammation? Why would the host rely on signals from foreign agents—adherent bacteria—to regulate vital defense decisions? Much remains to be understood about these and many other basic questions in the field [40–42].

In this manuscript we propose a minimal model of the dynamics at the mucus interface. The model in its simplest form is exactly solvable, affording the *in-silico* exploration of host strategies for sensing and responding to temporal changes in the intestine. Since living systems need time to adjust to change, we consider the effect of time delay in the host response and how delayed response can modify the optimal host sensing strategy. We outline the benefit the host derives from feed-forward control. We consider two types of signals: signals produced by *segmented filamentous bacteria* (SFB) which we consider as possibly anticipatory of the second type of signals, such as lipopolysaccharide (LPS), that are shed by the other bulk commensal microbiota. As a proof of concept, we use our model to integrate data from two recent publications, accounting for the defensive role of host-secreted antimicrobial peptides in the mouse small intestine. The modeling framework presented here is easily extendable to integrate further observations, and to consider situations where instead of being anticipatory the SFB convey other information, towards a deeper understanding of this essential system.

## RESULTS

### Model

Let us first think of this system rather generally, with microbiota derived signals and host-derived molecules to control the bacteria. A model of the system could be used to explore the benefits of various host defense strategies. Specifically, we suggest the role of feed-forward control in assisting the host to overcome inefficiencies due to response delays. Throughout the rest of this manuscript, we have a very specific realization of the biological system in mind, detailed below. But more broadly, the modeling framework we propose here is intended as a first step in developing a general theoretical framework for host-microbiota relations in the intestine. The key variables in our model are: the lumen microbiota and their products (LPS), special bacteria (SFB), and host derived molecules (antimicrobial peptides, AMP).

We focus on the ileum, a part of the small intestine where the dynamics of mucus defense are currently under intensive investigation. Here, the mucus does not self-organize to produce a dense impenetrable layer [4, 43]. Rather, the host defense relies on the secretion of AMP. Epithelial host cells detect bacterial proximity via microbial byproducts such as *lipopolysaccharide* (LPS) and *flagellin*. For the sake of simplification in our subsequent discussions, we will collectively refer to these microbial markers as LPS. These chemical cues are channeled through *MyD88* signaling [44] to sense bacterial proximity. When the bacteria are sensed to be close, or abundant, *MyD88* signaling leads to increased release of antimicrobial peptides (AMP) to the mucus environment. In healthy mice, this process results in the lumen microbiota being kept at homeostasis separated from the epithelial cells (Figure 1A). Thus, in the ileum, host-microbiota separation crucially relies on constant communication between the two systems. In *MyD88* – knockout mice, sensing of the LPS signals is significantly impaired, and as a result, the host cells do not excrete the AMP necessary to maintain the healthy separation [45]. The mucus loses its barrier properties and the bacteria encounter the host cells, causing inflammation and disease (Figure 1B). The full dynamics of the AMP-mucus system are complex, involving multiple AMPs secreted variably in the diurnal cycle. [30, 32, 46]

For a minimal model of the dynamics we focus on one important AMP, the *regenerating islet-derived protein 3γ*, Reg3g [24, 30, 32]. By first establishing this minimal modeling framework, more complex and realistic models may be developed thereafter as needed. To model the homeostasis at the barrier, which involves the constant sensing of bacterial products and secretion of antimicrobial peptides (AMPs), we propose a sense– and-secrete mechanism of conjugate diffusion: bacterial products such as *lipopolysaccharide* (LPS) and *flagellin*, as well as other products, are shed by and diffuse from the microbiota, reaching the host epithelium. Simultaneously, the AMP are produced by the host cells, and diffuse away from the epithelium towards the lumen. These dynamic processes feed-back on each other, forming the homeostatic state. Throughout this work, the coordinate *z* = 0 is the edge of the epithelium, and *z* increases until the lumen, perpendicularly away from the epithelial boundary. This is a much simplified geometry and one obvious avenue to expand our model, is to consider the influence of the anatomically correct structure, incorporating the spatial modulation of the villi, etc. In this manuscript, we elucidate the fundamental behavior of the system by concentrating on a simplified, one-dimensional representation of the mucin layer along the *z* axis (Figure 1C).

#### Model Equations

We collectively call the signals derived from the microbiota, disregarding the SFB, as LPS for convenience, with a concentration in space and time described by *c*(*z, t*). Similarly, the host derived antimicrobial proteins, (specifically, Reg3g), with concentration *a*(*z, t*). In our model, commensal (non-SFB) bacteria shed LPS [47, 48] that diffuses through the mucus and is detected by the epithelium (at *z* = 0); detection upregulates production and release of antimicrobial peptides (AMP) by the epithelium, which then diffuse through the mucus and inhibit the bacterial growth (Figure 1C). More than one bacterial species is present in the lumen microbiota. We define the local concentration of bacteria along the *z* axis by the vector (*z*) = {*b*_1_(*z*), *b*_2_(*z*), …}, for a set of bacterial species. Let the bacteria penetrate into the mucus (*z* decreasing) at velocities 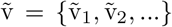. The AMP removes (kills) bacteria according to the vector of rates 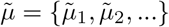, and LPS production rates *K* must be specified accordingly for each bacterial species. With these quantities defined, we may model the dynamics of the microbiota and host AMP. The non-SFB bacteria enter through the interface between the lumen and the mucus barrier, with dynamics 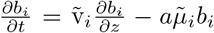. Together with the bacteria, bacterial products also diffuse in from the lumen-mucus interface as well as being produced by the bacteria present in the mucus layer. Therefore, the LPS dynamics read 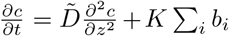. In response to sensing the bacterial products, the epithelial cells produce AMP, released at *z* = 0, and diffuse through the mucus, 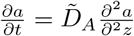. Here, 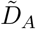 and 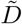 correspond to the AMP and LPS diffusion coefficients, respectively. We let the total mucus height be *L*_*M*_, and so, to reduce the parameter space, we introduce the following scaled quantities: 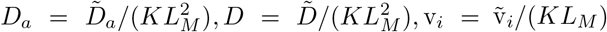, and 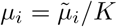,, where the index “*i* “ denotes the bacteria species. Furthermore, the distance from the epithelium is re-scaled, *z ← z/L*_*M*_ and *τ* = *tK*. In these scaled variables, the dynamics are:

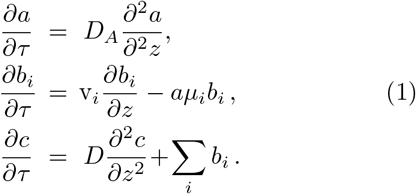

Now, *z ∈* [0, 1], with *z* = 0 corresponding to the boundary between the epithelium and the mucus, and *z* = 1 corresponding to the boundary between the mucus and the lumen. AMP is produced by the epithelium (*z* = 0) according to the sensed LPS, described in the *boundary conditions* section below. To account for AMP degradation upon activation on bacteria, one may add a degradation term to the AMP dynamics in Equation 1, e.g., −*a* Σ_*i*_ *μ*_*i*_*b*_*i*_. We found that the effect of degradation does not qualitatively change the results presented here (for a detailed comparison, see Appendix section A). Therefore, we focus on the simpler case, neglecting degradation.

#### Boundary conditions

To describe the host-microbiota interactions in the mucus, we must complement Equation 1 with biologically-meaningful boundary conditions. These include both lumen microbiota dynamics, and host regulation of AMP production. Deep inside the lumen, far from the mucus interface, the microbiota are understood to be largely unaffected by host AMP production [49]. We therefore make the simplifying assumption that at the mucus-lumen interface there are effectively no more AMP molecules [25, 50], therefore, *a*(*z* = 1) = 0 at all times. Furthermore, the lumen contains many non-SFB bacteria [24] and acts as a reservoir of their products, collectively called LPS in our model [51, 52]. Therefore, at the boundary between the mucus and the lumen we have *b*_*i*_(1) = *b*_*Mi*_, *c*(1) = *c*_*M*_, with *b*_*Mi*_, *c*_*M*_ the lumen bacteria and LPS concentrations, respectively. Furthermore, we assume that net LPS flux at the transition from lumen to mucus is zero at steady state, entailing 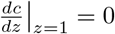

Finally, we must consider the AMP concentration at the epithelium-mucus interface, *z* = 0. This signifies the AMP production of the epithelial cells [53–55] and is an important factor the body must regulate. How should the AMP production be regulated? A trade-off exists between making a large enough amount of AMP to provide robust protection, but as little as possible. There are several reasons to minimize AMP production: (i) to save resources and avoid cell stress due to overproduction; (ii) Evidence shows that production of AMP in the wrong place can drive inflammation. It is possible that large amounts of AMPs produced in the ileum might find their way to the colon, where they could lead to generation of high levels of pathogen-associated molecular patterns (PAMPs) [56]. Indeed, appearance of Paneth cells in the colon, seen in ulcerative colitis, is thought to drive disease by generating these PAMPs [56, 57]. It is known that AMPs produced in the ileum are also present in the colon and feces [58, 59].

Although in principle there could be baseline production of AMP regardless of microbial product sensing, experiments show that if the sensing is stopped, so is AMP production [24]. Indeed, in germ-free mice, there is essentially no production of Reg3g, whereas other factors, such as lysozyme, are secreted regardless of colonization [60, 61]. Therefore, we ignored baseline production in our model of AMP. Our approach would apply similarly to other AMPs and microbial signals, though for these other cases perhaps a baseline production term would be suitable. Here, we initially consider *a*_0_ *≡ a*(*z* = 0) = *βc*(*z* = 0). Setting *a*(*z* = 0) as a boundary condition is equivalent to setting 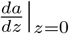, because setting its value at zero or its slope at zero are equivalent when it is pinned at *z* = 1 to *a*(1) = 0. For further discussion of the AMP boundary conditions *cf*. Appendix section F. The response-amplification constant, *β*, reflects the amplitude of the AMP response to the sensed LPS. To increase protection, the host would increase *β*. To save resources and reduce inflammatory load, the host would decrease *β*. Thus *β* controls the trade-off between defense and conservation.

Later, we will introduce a time delay in AMP response, between the time a signal is sensed (time *t − ϕ*_*a*_) and when AMP is secreted from the epithelium (time *t*). This delay will reflect the time taken to switch the transcriptional program, translate and assemble, and secrete the peptides. But, before examining the implications of delayed host response, we must first establish how mucus defense maintains host-microbiota equilibrium.

### AMP response to microbiota composition

If we assume the AMP are not degraded we can solve Equation 1 at steady state. Letting 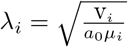 we have,

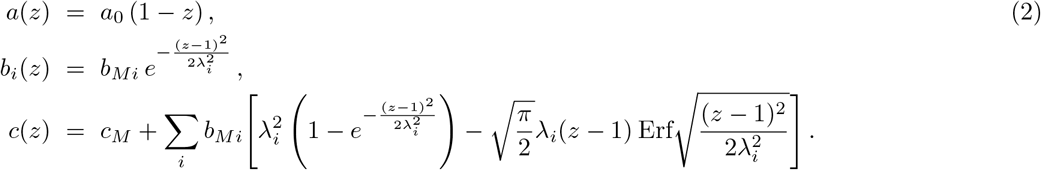

where Erf stands for the error function and *λ*_*i*_ defines the effective penetration length of the bacteria from the mucus-lumen boundary at *z* = 1. The bacteria penetrate from *z* = 1 according to a Gaussian shape, while the AMP diffuse from *z* = 0 forming a linear gradient of slope *a*_0_. The bacteria have an additive effect through their LPS shedding, evident in the sum in the solution for *c*(*z*). A more detailed examination of this effect is explored in Appendix section C.

Figure 1D illustrates the local concentrations of AMP and of a single bacterial species as function of the distance (*z*) from the gut-microbiome interface for *b*(*z* = 1) = *b*_*M*1_ = 1, *c*_*M*_ = 1. In our focus on a single species model, *a*_0_ is a control parameter for *λ*_*i*_. Therefore, we may set v_*i*_*/μ*_*i*_ = 1 without sacrificing the generality of our findings.

As the experimentally observed bacterial profiles can generally be scaled by a bulk value, *b*_*M*_, and the distance *z* is also already scaled, by extracting *λ*_*i*_ from imaging data one would be able to characterize the system.

### Feed-forward control primes the host to changes in the bacterial background

There may be several mechanisms by which a host may gain pre-warning of an incipient rise in lumen bacterial levels. Such feed-forward control would be especially important if the host response is not immediate, but rather, delayed. We proceed to consider feed-forward AMP production as a response to the SFB signals, assuming these SFB signals anticipate the lumen microbiota levels.

Recent experiments carried out by Brooks et al. [30] suggest that feeding promotes rhythms in the epithelial attachment of SFB, stimulating the expression of AMP. The SFB appear to be harmless and are interpreted as a signal, rather than as antagonists that necessitate a defensive response. Indeed, in a different system, the SFB are essential for regulating the levels of mucosal T-helper 17 cells [62]. But why would the host use SFB signals? We consider the effect of cyclic diurnal changes.

To accommodate the cyclic diurnal changes, we incorporated two temporal delay parameters into our model: *ϕ*_*a*_ for the delay in AMP production, and *ϕ*_*b*_ for the delay in microbiota growth. The time delays *ϕ*_*a*_ and *ϕ*_*b*_ are measured with respect to the time of SFB translocation to the epithelium; one possible mechanism for SFB translocation is by the nutrient flux generated when food arrives to the lumen [30]. Conversely, the bulk lumen microbiota experience abundance changes rather than translocation, therefore operating with a different time lag. Hence, we considered scenarios where the SFB translocate and stimulate the epithelium earlier than the bulk lumen bacteria rise. Measuring time since the SFB arrival: *ϕ*_*a*_ estimates the duration from epithelial detection, signal processing, transcription factor modification, to AMP assembly and secretion; *ϕ*_*b*_ represents the response time of the lumen microbiota, and accordingly their LPS signal, to nutrient arrival. Being able to investigate host strategies for various *ϕ*_*a*_, *ϕ*_*b*_ combinations *in silico* may be useful to generate experimental hypotheses.

To prioritize the sensing between the two distinct signals, SFB and LPS (from the non-SFB microbiota), we introduce the parameter *ν*. Accounting for the time delays in AMP and bacteria, we update the boundary conditions. The lumen microbiota rise and ebb, and the AMP is stimulated by two channels, with

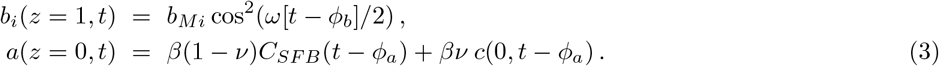

The host, for AMP production, can sense both LPS (*ν* = 1) and SFB (*ν* = 0) signals. Our study clarifies the advantage that this dual-channel sensing confers to the host.

Phenomenologically, the SFB translocate to the epithelium according to *C*_*SFB*_(*t*) = *b*_*Mi*_ cos^2^(*ωt/*2), while *c*(*z* = 0, *t*) represents the concentration of LPS-like molecules to which epithelial cells are exposed at time Note that the SFB and LPS signals have the same dynamical range, *b*_*Mi*_, when no AMP is present (*β →* 0). When fitting the data, *ω* should be thought of as a phenomenological parameter for the 24-hour time-series given—the cos^2^(…) form represents a minimal theory and not necessarily the true dynamics—current experimental measurements are not at sufficiently high temporal resolution to suggest more realistic models of the dynamics. As higher resolution time-series become available, it would be easy to amend these boundary conditions to be more realistic. Importantly, in what follows, in this work we assume *quasi steady-state* meaning that the AMP, bacteria, and LPS remain at steady state values and their temporal changes only reflect the changing boundary conditions and built-in time delays. A simple diffusion-based argument justifies the quasi steadystate assumption: whereas the boundaries (lumen, AMP production) change on the scale of 4-6 hours, an AMP molecule with a diffusion coefficient of 10^*−*6^*cm*^2^*/s* would require approximately 100 seconds to traverse the 100*μm* wide mucus layer [63]. This separation of time-scales permits us to treat all diffusion as at steady-state, even as the boundaries slowly pendulate.

How do varying, sometimes conflicting signals from the SFB and microbiota affect host AMP production? Figure 2 displays the concentration of bacteria, AMP, and SFB as a function of time; the different panels show different values of *ν*—the relative weight in sensing SFB vs. LPS. For the sake of comparison, we also present the boundary condition *b*_1_(*z* = 1, *t*) = *b*_*M*1_ cos^2^(*ω*[*t − ϕ*_*b*_]*/*2). Mimicking the dynamics measured in Ref. [32], and studied later in this manuscript, we picked experimentally-relevant parameters, with *ϕ*_*a*_ = 3 and *ϕ*_*b*_ = 1 (Figure 2). The left (*ν* = 0) and right (*ν* = 1) panels of Figure 2 represent, respectively, the scenarios where the AMP response is driven either solely by the SFB or, conversely, only by the LPS produced by the bulk microbiota. Upon increasing from *ν* = 0 to *ν* = 0.25, we observe a reduction in the total amount of AMP (blue shaded area) with an increase in bacterial exposure (red shaded area). Continuing to *ν* = 0.5 the further increase in bacterial exposure may exceed the acceptable threshold. Thus, an optimal solution that balances good defense with limited resource expenditure would be around *ν* = 0.25. The results shown in Figure 2, however, are for a specific combination of time delays. We were therefore motivated to clarify optimal host behavior in terms of our model parameters: strength of response, *β*, and focus of attention, *ν*.

**FIG. 2.**
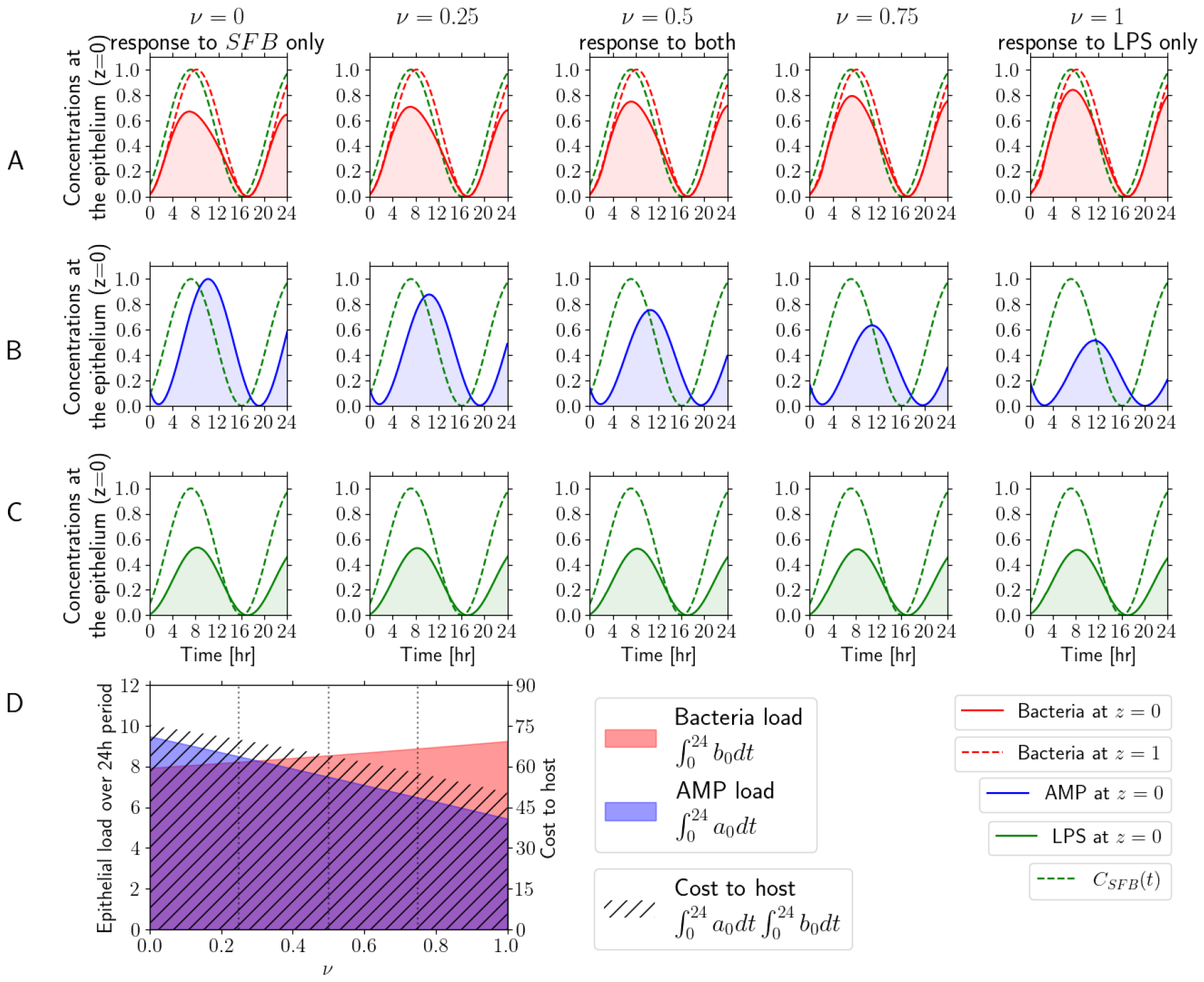
Optimal host defense depends on division of attention between the SFB and LPS signals. (A-C) Concentrations at the epithelium (*z* = 0) as function of time, in hours, for bacteria (in red, panels in row A), AMP (in blue, panels in row B), and LPS (in green, panels in row C). We take *δt* as a universal time-shift that defines what hour ‘zero’ is. The dashed red lines mark the lumen microbiota concentration *b*(*z* = 1, *t*) = *b*_*M*1_ cos^2^[*ω*(*t −ϕ*_*b*_ *− δt*])*/*2], with a *ϕ*_*b*_ delay. The dashed green line is the SFB signal *C*_*SFB*_ (*t*) = *b*_*M* 1_ cos^2^[*ω*(*t − δt*)*/*2] which is without delay. Each column corresponds to a different value of *ν*. Left: the AMP responds only the SFB signal, ignoring the LPS (*ν* = 0); Middle: sense both LPS and SFB equally (*ν* = 0.5). Right: sense only the LPS signal, ignoring the SFB (*ν* = 1). Mimicking the dynamics measured in Ref. [32], studied later in this manuscript, we set *b*_*M*1_ = 1, *ϕ*_*a*_ = 3, *ϕ*_*b*_ = 1; *δt* = 7.2 hours and *ω* = *π/*8.9 hours^*−*1^, *β* = 1. (D) Epithelial load over 24 hours, as function of *ν*. The bacteria load in red: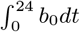 in blue: AMP load 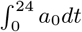 hatched: cost to host 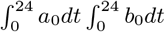. All results were obtained for *b*_*M* 1_ = 1, *D* = *D*_*A*_ = 1, and v_1_*/μ*_1_ = 1.

### Combined feed-forward and ambient sensing of SFB and LPS can be used to optimize host response

The choices *ν* = 0 or *ν* = 1 represent two extreme cases: response induced only by SFB or only LPS signals, respectively. Given that both signals potentially convey useful information about the environment, there may be an advantage to be gained from responding to both signals—therefore, an intermediate value of *ν* may be optimal. Yet, optimal in what sense? Maximal defense involves minimizing exposure to bacteria, therefore, at the epithelium surface (*z* = 0) one goal is to minimize 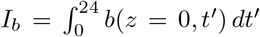. However, this goal is satisfied when *b* = 0, which requires an infinite amount of AMP. Instead, suppose that the epithelium tolerates a specified maximal amount of bacteria in a 24-hour period without inflammatory changes. To minimize the host cost, consider 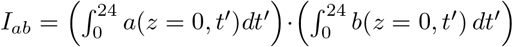, requiring that the exposure (*b*) does not exceed the maximal tolerated load. The cost to the host is deemed proportional to the amount of AMP produced (Figure 2C). We considered alternative cost functions and their effect, and concluded that the results presented in this manuscript do not depend qualitatively on this specific form of the cost function (*cf*. Appendix section E).

There is fitness pressure on the host to tune *ν* and *β* to optimize defense: minimizing defense cost while maintaining the required protection target. We therefore define the optimal response parameters, *ν* and *β*, such that the bacterial exposure ∫ *b*(0, *t*^*′*^) *dt*^*′*^ is the tolerated exposure, but as little as possible AMP is produced, thus preserving resources. We explore the optimization landscape by producing heatmaps for two specific choices of *ϕ*_*a*_ and *ϕ*_*b*_ in Figure 3. For a given pair, *ϕ*_*a*_ and *ϕ*_*b*_, there exist a pair, *ν* and *β*, that minimize AMP production while not exceeding the bacterial exposure threshold (white dashed contours in Figure 3). This optimal point is shown as the white dot in the heatmaps in Figure 3. Where *ϕ*_*a*_ = *ϕ*_*b*_ = 1, we observe that the minima are always located at *ν* = 0, this is not surprising. Essentially, the SFB, which do no have a time delay in this description, anticipate the bacteria (delay *ϕ*_*b*_) perfectly in concert with the host AMP production delay (*ϕ*_*a*_), so the host minimizes resource expenditure by listening solely to the SFB signal. However, when *ϕ*_*b*_ is much larger, a more complex behavior emerges: the mismatch between *ϕ*_*a*_ and *ϕ*_*b*_ yields minima that progressively move towards larger values of *ν* upon decreasing *β*; for example, the minimum value of the uppermost line 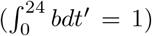 moves from *ν* = 0 when *ϕ*_*a*_ = *ϕ*_*b*_ = 1 to *ν ≈* 0.5 for *ϕ*_*a*_ = 1 and *ϕ*_*b*_ = 3. The model thus encapsulates optimal scenarios that monitor either a single channel (only LPS or only SFB) or both channels simultaneously.

**FIG. 3.**
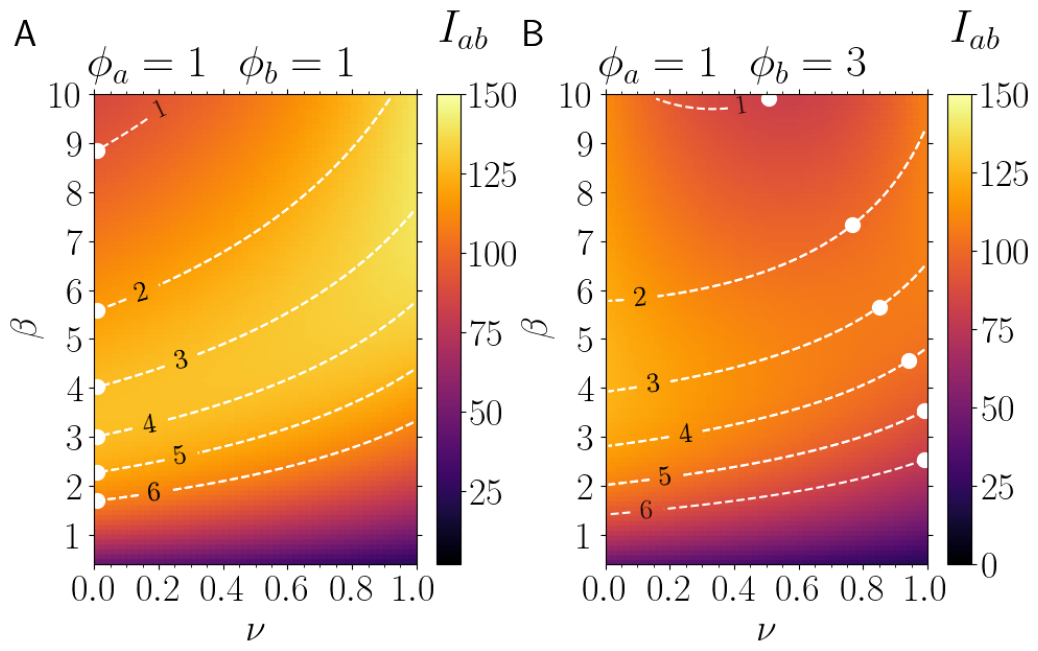
Finding the optimal host response. Heatmaps for *β* versus *ν* showing optimal host defense. For a fixed bacterial load—dashed lines indicate equal-contour lines, 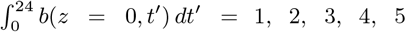 and 6, from top to bottom—the minimal host cost in our model is the *β* and *ν* pair on this contour that minimize the cost 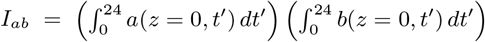, shown as a white dot. (A) *ϕ*_*a*_ = 1, *ϕ*_*b*_ = 1, the SFB anticipate the LPS perfectly, the optimal *ν* is zero. (B) *ϕ*_*a*_ = 1, *ϕ*_*b*_ = 3, the lumen microbiota lag significantly behind the SFB, the optimal *ν* involves listening to both channels, 0 *< ν <* 1. Results for *δt* = 7.2 hours, *ω* = *π/*8.9 hours^*−*1^, *b*_*M* 1_ = 1, *D* = *D*_*A*_ = 1, and v_1_*/μ*_1_ = 1. Further information about the bacterial load can be found in the Appendix section B.

The microbiota growth delay, *ϕ*_*b*_, when compared to the SFB, can vary depending on diet, microbiota composition, location in the intestine, and so forth. Similarly, host cell AMP production delay, *ϕ*_*a*_, may vary significantly with respect to host regulation and cell-to-cell variability. Therefore, what are the optimal *ν* and *β* values for all combinations of *ϕ*_*a*_ and *ϕ*_*b*_? Maintaining fixed bacterial exposure in a 24 hour period, here, 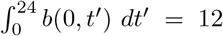, these optimal *ν* and *β* pairs are shown in Figure 4. As we maintain a fixed value for *I*_*b*_ = *b*(0, *t*^*′*^)*dt*^*′*^, the optimal *β* shown in Figure 4B are proportional to the total cost to the host. In the limit where the AMP production responds very quickly upon receiving the signals—indicated by a small *ϕ*_*a*_—the optimal defense strategy imposes minimal costs. Similarly, the cost is small when the bacterial load is also delayed— indicated by large *ϕ*_*b*_. Thereby our model captures the pressure on the host to develop SFB-like mechanisms to optimize response. Futhermore, the assymetric nature of the cost with regards to *ϕ*_*a*_ suggests pressure on the host to produce AMP as quickly as possible or face a large cost.

**FIG. 4.**
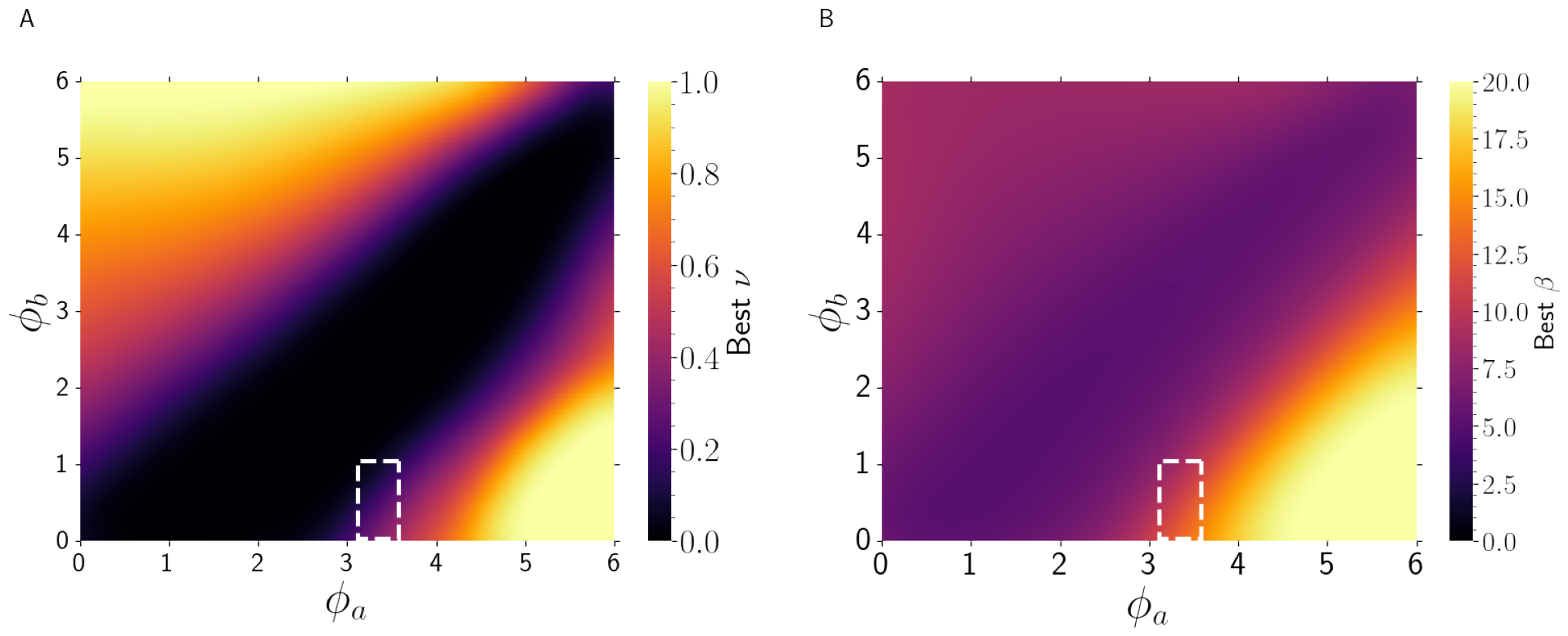
The optimal defense hypothesis, assuming the SFB anticipate the LPS signal. Heat-maps for *ϕ*_*a*_ versus *ϕ*_*b*_ color-coded according to the optimal values of *ν* (left) and *β* (right). The optimal *ν* and *β* values correspond to the coordinates of the minimum points (i.e., the white dots in Figure 3) along the contour 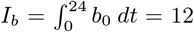. In both diagrams, the white-dashed region marks the range of parameters *ϕ*_*a*_ and *ϕ*_*b*_ specified by the fits in Figure 5 and discussed in the application to data section of this manuscript, below. All results were obtained for *b*_*M* 1_ = 1, *D* = *D*_*A*_ = 1, and v_1_*/μ*_1_ = 1.

The fact that we obtain optimal values of *ν* that are not 0 or 1 is interesting, because it means that within the model assumptions, there are regimes where host behavior benefits from integrating both signals. The existence of an optimal strategy suggests that there is evolutionary pressure towards this optimum, we therefore build on this notion and hypothesize that host defense is regulated towards such optimal behavior. We proceed to apply our model to published data, presenting evidence supporting the optimal defense hypothesis.

### Proof-of-concept application of the model to circadian AMP measurements

Our model can be applied directly to experimentally observed rhythms in intestinal bacteria and AMP. As a proof of concept, we turned to experimental data reported by Brooks *et al*. [30] and Frazier et al. [32] and fitted our model to these data (Figure 5). The model receives as inputs the parameters *ϕ*_*a*_, *ϕ*_*b*_, *β, ν, ω* and *δt*, which are obtained by fitting the experimental data; the Appendix section D contains further details about the fitting procedure and data normalization. Strikingly, we could place both Frazier’s and Brooks’ AMP measurements, two independent experiments by different research groups, on a single curve without specific tuning (Figure 5C).

**FIG. 5.**
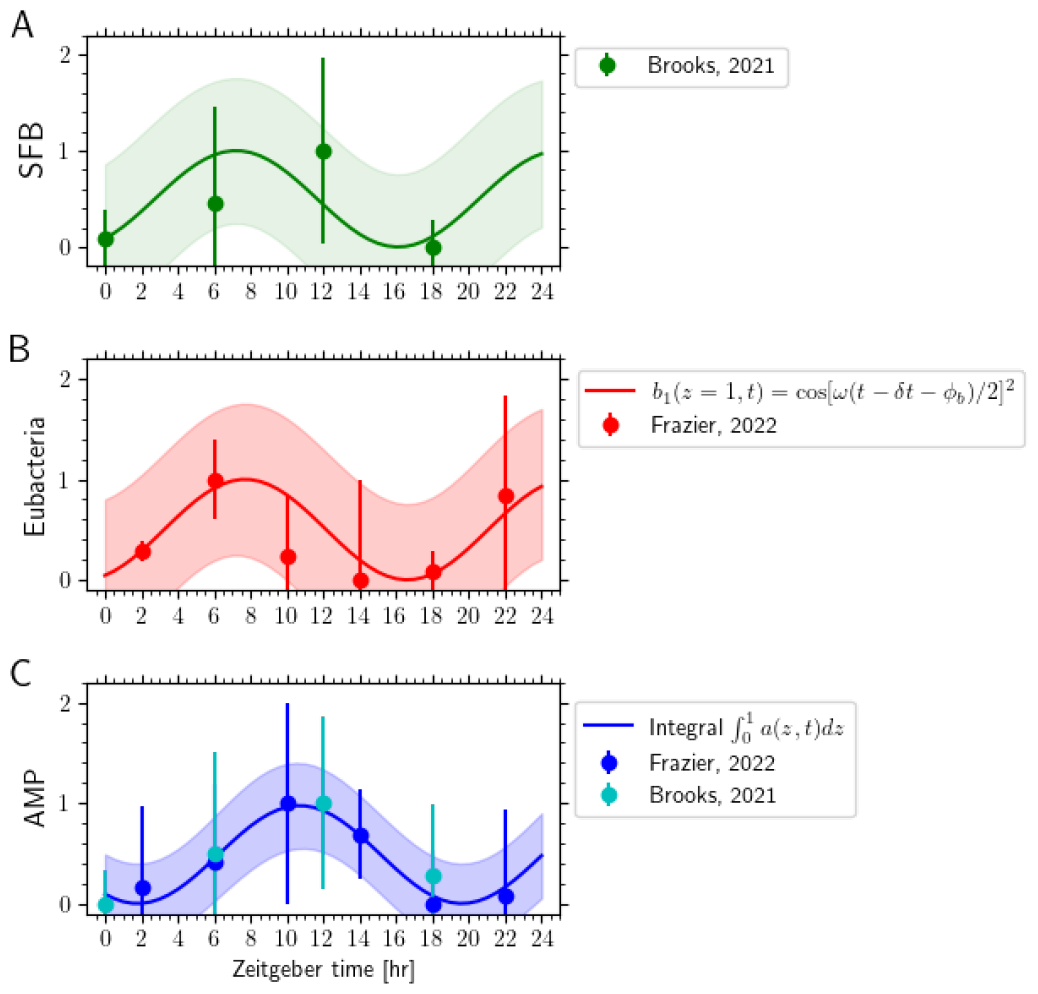
Proof-of-concept application on experimental data. Comparison between model predictions (solid curves) and experimental data previously reported [30, 32] according to the legend. (A) SFB attached to the epithelium, measured by Brooks *et al*.. (B) 16S rRNA gene amplicon sequencing of distal ileum luminal. (C) AMP (Reg3g) quantified using immunostaining fluorescence intensity (Frazier) and immunoblot densitometry (Brooks). Fitting parameters: *δt* = 7.20 *±* 0.59, *ϕ*_*a*_ = 3.30 *±* 0.20, *ϕ*_*b*_ = 0.50 *±* 0.66, *w* = *π/*8.90 *±* 0.32, *β* = 2.46 *±* 0.87, and *ν* = 0.30 *±* 0.37 (all to 95% confidence). The shaded regions in each diagram define the 95% fit confidence interval. All results were obtained for *b*_*M* 1_ = 1, *D* = *D*_*A*_ = 1, and v_1_*/μ*_1_ = 1.

Fitting the timeseries gives *ϕ*_*a*_ = 3.49 *±* 0.19 and *ϕ*_*b*_ = 0.50 *±* 0.66 hours, *β* = 2.83 *±* 0.01, and *ν* = 0.31 *±* 0.05, (results to 95% confidence). Accordingly, both SFB and LPS channels matter for AMP production, consistent with current knowledge. We turned to Figure 4 to see if the fit results are consistent with the optimal defense hypothesis we proposed. The values of *ϕ*_*a*_ *≈* 3.5 and *ϕ*_*b*_ *≈* 0.5 are depicted in the dashed-white boxes in Figure 4. With regards to optimal *ν*, these values are consistent with the fit *ν ≈* 0.3 (purple color). Interestingly, both *ϕ*_*a*_ and *ϕ*_*b*_ seem to sit at the boundary between the region where the optimal *ν* is the single-channel *ν* = 0 and where listening to both channels is worthwhile. We consider possible implications of being on this boundary in the Discussion. With regards to agreement with *β* for the optimal defense hypothesis, here some mismatch occurs. However, whereas with *ν*, having a value which is not 0 or 1 is qualitatively different since it suggests the role of both channels in the process, when it comes to *β*, there is not such qualitative test, apart from trivially *β >* 0. Therefore, the true value of *β* requires specific details outside our model. Nevertheless, we note that the optimal defense (Figure 4) suggests that the value of *β* is at the boundary between the low *β* and high *β* regions.

## DISCUSSION AND CONCLUSIONS

In this study, we developed a model of the host’s antimicrobial defense in the ileum, and how it prevents the invasion of the lumen microbiota through the mucus barrier. Despite considerable experimental focus on this system, holding potential for significant medical advancements, this study, to our knowledge, presents the inaugural mathematical model for it. Here, we proposed a conjugate diffusion model in which the host detects the presence of microbes and responds by secreting AMP (Figure 1). Using the model, one can integrate experimental observations into a single framework, enabling testing of hypotheses against model predictions and guiding experimental efforts.

Describing the system, our model solution depends on a bacterial penetration length, denoted by *λ*, which represents the fraction of the mucus layer that bacteria from the lumen can successfully invade at steady state. Extracting this length from imaging data is theoretically feasible, but technically challenging, because imagining the mucus in the ileum is difficult. One possible approach is to use 16S fluorescent in situ hybridization [24]. Although in a natural community we would expect the values of *λ*_*i*_ to differ between species, an effective (phenomenological) community penetration depth could still be used for a coarse-grained description of the system. Moreover, we demonstrated that under certain simplifying assumptions, the host defense mechanism is insensitive to the exact composition of the lumen microbiota, and instead, reacts to their total abundance, thus justifying the use of a single, community-level *λ*. As needed, our minimal framework could be expanded to include more complex and realistic features.

The abundance of the intestinal microbiota oscillates during the diurnal cycle, in tune with feeding behavior. Hence, optimal defense of the mucus barrier requires synchronization of AMP secretion with the diurnal microbial cycles. But production of AMP takes time. Let us assume that the signaling cascade triggered by the cellular sensing of microbial ligands takes 15 minutes to reach steady state. To secrete AMP, the cells must then undergo transcription, translation, assembly, and secretion, a process that can take from 15 minutes to several hours. How does the delay in host AMP secretion shape the host’s sensing strategy?

Given delayed host response, a feed-forward control mechanism could be used to optimize host response. While there may be several ways the host can get pre-warning of an incipient increase in lumen bacterial levels, in this manuscript we considered a specific biological system. Based on recent experimental understanding, we introduced a second source of stimulus in our model: *segmented filamentous bacteria* (SFB), which are known to stimulate the host AMP diurnal cycle. If the SFB anticipate the lumen microbiota, by simultaneously monitoring the LPS and SFB signaling pathways, the host can respond to the current environmental conditions (LPS) and anticipate future changes (SFB). Conversely, if the SFB do not anticipate the lumen microbiota, they may still encode important information: the SFB have privileged access to signaling pathways due to their unique capability to adhere to the epithelium. We quantified the host attention using *ν*, the fraction of AMP response that is attributable to the LPS vs. SFB pathways (Figure 2). As with any defense mechanism, there is a cost associated with the host response to microbial invasion. In this study, we elucidated how the host may minimize defense costs while sustaining the necessary level of protection. We explored alternative cost functions, all yielding qualitatively similar behavior (*cf*. Appendix section E).

The host must avoid excess killing of the microbiota, else it would risk being flooded with inflammatory signals such as pathogen-associated molecular patterns (PAMPs) [56]. One way for the host to optimize its response is by limiting the total bacterial load at the epithelium boundary over 24 hours while minimizing AMP production (Figure 3). We propose an ‘optimal defense hypothesis’, wherein the host minimizes defense cost by identifying the ideal *ν* value that maintains bacterial load at the epithelial boundary within a specified 24-hour threshold. We explored this optimization strategy for various time delays both in AMP production and in microbiota growth (Figure 4). If the SFB anticipate the lumen microbiota, then the host may benefit from considering the SFB signal. In some scenarios, the model suggests that optimal defense involves a response to both LPS and SFB signals, with 0 *< ν <* 1. This contrasts with the notion of exclusively responding to a single signal. When we applied our model to analyze data from two recent publications, it was able to fit both AMP (Reg3g) measurements reported by different research groups onto a single curve (Figure 5). The fitting results suggest that host AMP response delay takes about 3 hours after stimulus, whereas the lumen microbiota exhibits little lag following the SFB. The host’s ability to monitor both stimuli (*ν ≈* 0.3) is consistent with the optimal defense hypothesis.

Although the fit results presented here demonstrate how to apply our model to experimental measurements, they were based on a limited number of time-points gathered from the literature. Therefore, caution must be taken when interpreting the results, and further data is needed to develop the model. Specifically, the phenomenological cos^2^(*ωt*) should be improved. We hope that this study will inspire researchers to collect more densely-sampled time-series data in order to enable more sophisticated analyses of this crucial system.

The assumption made in this study, that the lumen microbiota are equally susceptible to AMP, precludes pathogens known to be resistant to certain AMP [30]. To model such pathogens with different susceptibilities to antimicrobial peptides, each pathogen would be assigned a distinct *λ*_*i*_ value, reflecting its individual resistance level. A deeper study would consider not only the AMP resistance, but also the capacity of the pathogens to swim along and inside the mucus and to attach to the epithelial cells [64]. The use of specific rRNA probes could enable imaging and quantification of such pathogens. One may also use mutant bacteria, lacking certain swimming capabilities [65]. Moreover, by using drugs that stimulate or block the production of AMP it may be possible to change *ϕ*_*a*_ and see if the host responds with altered *β, ν*, dynamically.

When formulating the optimal defense hypothesis presented in this work, we assume that the SFB signal anticipates the lumen microbiota, for there to be an advantage to listen to the SFB. This assumption may be relaxed to a simpler assumption: that the SFB add information to what could be learned from the signals emitted by the lumen microbiota alone. Instead of being anticipatory, conveying information about the time domain, one may consider how SFB contribute to host sensing by activating alternative signaling pathways, thereby making host sensing more robust through integrating different inputs.

Why might the host evolve reliance on SFB as part of an optimized AMP defense strategy? Relying on external factors for a critical defense mechanism can be perceived as risky. Here, we demonstrated that feed-forward control combined with feedback from the ambient microbial background confers an advantage. While SFB is the feed-forward signal we considered in our study, it is possible that the host’s nervous system could trigger AMP production as it sends hunger and feeding signals, thus avoiding dependence on other organisms. So, why does the host rely on SFB instead of solely triggering AMP production in response to feeding-induced signals sent by nerves?

One potential explanation is that the ability to sense microbial background developed earlier in evolution, and with it the capacity to use SFB-like microbes for feed-forward sensing. Indeed, it is possible that an SFB mechanism for immune regulation may have already been operational when the AMP system evolved [62]. After fulfilling the need for feed-forward sensing through SFB-like microbes, it’s likely that further evolutionary pressure for alternative solutions was insufficient.

An alternative explanation relies on the effect of fasting on the SFB signal. As recently demonstrated by Brooks et al. [30], no SFBs were visible under fasting conditions. This provides an additional control mechanism: under fasting conditions, the likelihood of the next meal decreases, and with it, the effect of the SFB signal. Response to fasting prevents the unnecessary expenditure of AMP resources during periods of nutrient scarcity. Therefore, reliance on SFB signaling, which is responsive to fasting conditions, may offer an advantage to the host during uncertain meal timings.

By enriching our model and using denser time-series data, we envisage substantial advancements in deciphering the dynamics of AMPs. Ultimately, by uncovering the general principles governing mucus defense mechanisms, we may learn ways to modulate them, leading to potential cures for related disorders. A notion we explored here—optimal defense—may be one of these general principles. While our model of the mucus AMP defense system is minimal, it provides a testable framework that aligns with current empirical observations, supporting optimal defense. Therefore, we propose that analogous SFB-like mechanisms may benefit other species, including humans, and apply to mucus barriers other than in the ileum.

## ACKNOWLEDGMENTS

The authors acknowledge the Hebrew University of Jerusalem for the financial support and computational resources. The authors are also grateful to Dr. J. Brooks for kindly providing the experimental data used in Figure 5. G. V. B. also acknowledges Vicerrectoría de Investigación, Desarrollo y Creación Artística (VIDCA/UACh Project INS-INV 2022-09) for financial support.

All code and data can be found in https://github.com/AmirErez/GutConjugateDiffusion.

## SI Appendix

### Appendix A: Consumption of AMP when killing bacteria

One might think of a situation where AMP’s are degraded by the bacteriocidal process. In Appendix A— figure A we display the results for the local distributions of AMP, bacteria, and LPS obtained in the scenarios with and without AMP degradation. For the sake of a better resolution, we have set *b*(*z* = 1) = *b*_*M*_ = 5.5 and *c*(*z* = 1) = *C*_*M*_ = 2*b*_*M*_ for the boundary conditions.

We observe that the AMP degradation has a slightly more pronounced impact on the bacterial concentration for smaller *β*, with a larger number of bacteria being able to reach the epithelium at *z* = 0. For larger *β*, however, the AMP response is strong enough that practically no bacteria can reach the interface at *z* = 0. Since the overall differences between both scenarios are minor, throughout this study we have assumed 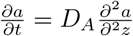 for the sake of mathematical simplicity.

**FIG. S1.**
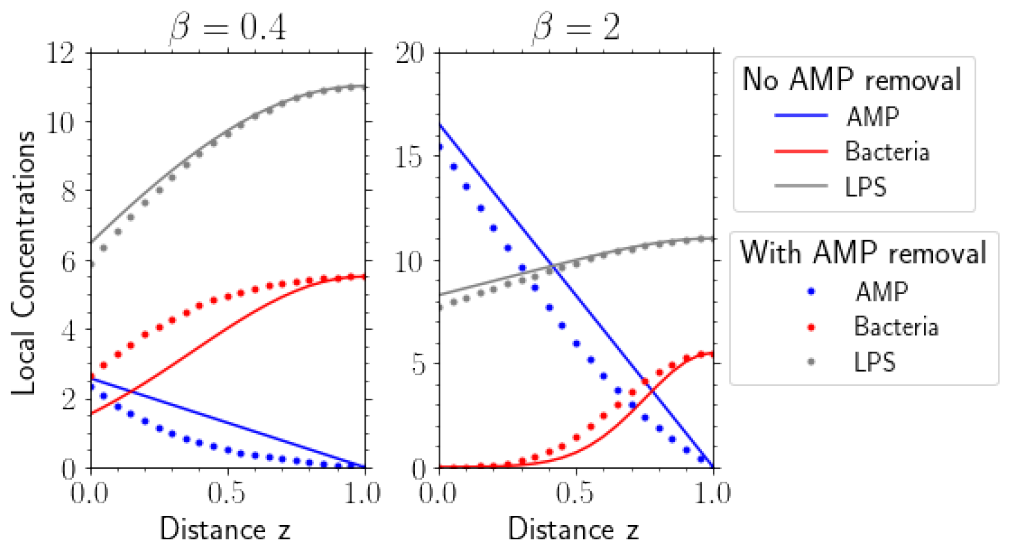
Concentration profiles. Local concentration as function of the scaled distance z away from the gutmicrobiome interface for *β* = 0.4 (left) and *β* = 2 (right). In gray, blue, and red are displayed the value of LPS, AMP, and bacteria, respectively. Solid curves correspond to results without AMP consumption whereas the color-matching symbols denote the values obtained when AMP is consumed. In all cases, the bulk concentration of bacteria is *b*_*M* 1_ = *b*(*z* = 1) = 5.5. All results were obtained *D* = *D*_*A*_ = 1, and v_1_*/μ*_1_ = 1.

### Appendix B: Bacterial Load

The optimal host response was analyzed in Figure 3 by means of *β* versus *ν* heatmaps for the integrals 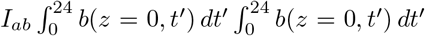. There, we have used symbols to mark the minimum value that such integrals assume along the contour-lines corresponding to the bacteria load, i.e.,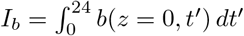. It is noteworthy, however, that the set of *β* and *ν* that minimizes the bacteria load does not necessarily correspond to the minimum in the host response. In order to illustrate that, in Appendix B—figure S2 we present how the bacteria load varies as function of *β* and *ν* for *ϕ*_*a*_ = *ϕ*_*b*_ = 1h (panel A), *ϕ*_*a*_ = 1h and *ϕ*_*b*_ = 3h (panel B), and *ϕ*_*a*_ = 1h and *ϕ*_*b*_ = 6h (panel C).

**FIG. S2.**
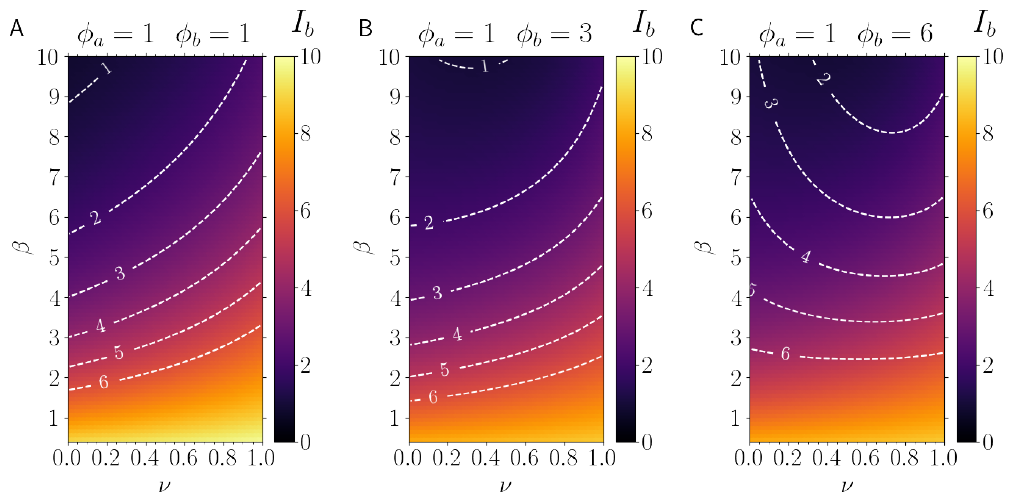
Total bacterial load in a 24-hour period. Heatmaps of *β* versus *ν* color-coded according to the bacterial load. with *ϕ*_*a*_ = *ϕ*_*b*_ = 1h (panel A), *ϕ*_*a*_ = 1h and *ϕ*_*b*_ = 3h (panel B), and *ϕ*_*a*_ = 1h and *ϕ*_*b*_ = 6h (panel C). For a fixed bacterial load (dashed lines indicate equal-contour lines), 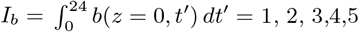, and 6, from topto bottom as indicated. All results were obtained for *b*_*M* 1_ = 1, *D* = *D*_*A*_ = 1, v_1_*/μ*_1_ = 1, *δt* = 7.2 hours, and *ω* = *π/*8.9 hours^*−*1^.

In panel A, where the response is in sync with the bacteria, *ϕ*_*a*_ = *ϕ*_*b*_ = 1h, we note that all the minima occur at *ν ≈* 0, listen to SFB only. In panels B, and even more pronounced in C, the asymmetry in *ϕ*_*a*_ and *ϕ*_*b*_ pushes the minima towards larger values of *ν* –listen also to LPS.

### Appendix C: Additive effect of different bacteria with the same properties

The present model allows us to investigate scenarios where different bacteria species reach different values at the mucus boundary, *z* = 1. Let us denote by *b*_*Mi*_ as the concentration of bacteria of species *i* at the position *z* = 1, i.e., *b*_*M*1_ = *b*_*i*_(*z* = 1). In Appendix C—figure S3 we display the total local concentration of bacteria of types 1 and 2, that is, *b*_1_(*z*) + *b*_2_(*z*), as function of the distance *z* away from the ileum-microbiome interface. The left panel is for the case where both bacteria types have the same bulk value; for the sake of a better visualization, we set *b*_*M*1_ = *b*_*M*2_ = 5.5 and all results are displayed in blue color. The right panel, conversely, corresponds to the scenario where *b*_*M*1_ = 1 and *b*_*M*2_ = 10, with results shown in red; note that in both cases, the total amount at the mucus boundary is the same,i.e. *b*_*M*1_ + *b*_*M*2_ = 11. We then note that, for a fixed value *b*_*M*1_ + *b*_*M*2_ = 11, the total amount of bacteria at any position *z* is the same irrespective of the individual values of *b*_*M*1_ and *b*_*M*2_. Such an statement is also valid when we compare the corresponding results obtained with and without AMP removal.

**FIG. S3.**
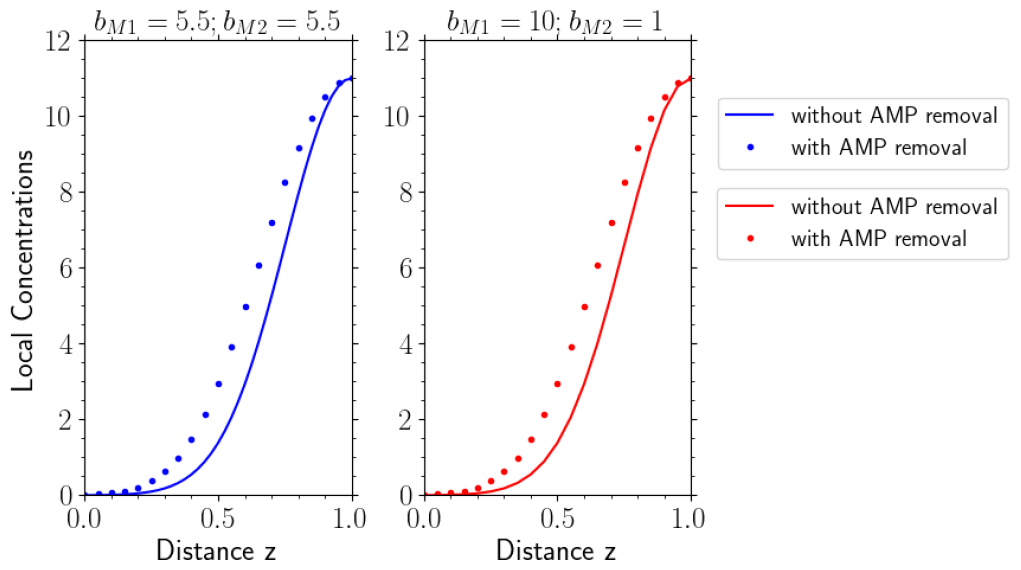
Two species of bacteria. Total amount of bacteria of two bacteria species at a position *x* for the scenarios with and without AMP consumption, shown as symbols and color-matching solid lines, respectively. Left diagram is the results for equal bulk values for both bacteria species, *b*_*M* 1_ = *b*_*M* 2_ = 5.5. In the right diagram one bacteria has a bulk value *b*_*M* 1_ = 10 and the other *b*_*M* 2_ = 1. Results obtained for *D* = *D*_*A*_ = 1, and v_1_*/μ*_1_ = v_2_*/μ*_2_ = 1.

### Appendix D: Fitting the Model to Experimental Data

In Figure 5 we display how our model predictions compare with experimental data reported by Refs. [30] and [32]. In order to obtain these results (solid curves), the model parameters were determined via the following fitting procedure:

1. All the experimental data – bacteria (namely, mucus-associated eubacteria in wild-type mice), AMP and SFB – are normalized to be in the range from 0 (smallest value) to 1 (largest value); such a normalization applies to both the observed values and to the corresponding errors. Hence, we have *b*_*M*1_ = 1 and, as previously, also set *v*_1_*/μ*_1_ = 1. The normalized data then constitutes the data set used in the next steps.
2. The SFB and eubacteria values are fitted in the same procedure, via the function *NonLinearModelFit* – from *Wolfram Mathematica v*.*12* – with the fitting parameters *w, ϕ*_*b*_, and *δt*.
3. With the three parameters above determined, we recall the boundary condition *a*(*z* = 0, *t*) = *β*(1 *− ν*)*C*_*SF B*_(*t − ϕ*_*a*_) + *βν c*(0, *t − ϕ*_*a*_) and use the *NonLinearModelFit* and *FindRoot* functions to self-consistently fit the AMP data in terms of the remaining last three parameters: *β, ν*, and *ϕ*_*a*_.
4. In all steps, the errors associated with the parameters estimates are listed by the command *NonLinearModelFit[“ParameterTable”]*.
5. We have used Frazier’s measurements for the Eubacteria because they have more timepoints and lower variance.

### Appendix E: Alternative cost functions

Do the conclusions we arrived at in the manuscript depend on a specific choice of the cost function? To answer this question, we investigated the effect of alternative cost functions. First, instead of following the total bacterial load over 24 hours, i.e. ∫ *b*(*z* = 0, *t*) *dt*, we focus on the maximum epithelial concentration of bacteria over the 24 hours, 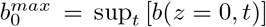. The logic behind such a cost function is that the host may need to defend against a temporary spike in bacteria instead of the total exposure over the 24 hours. In Figure S4 we present the heatmaps for *β* versus *ν* showing optimal host defense, 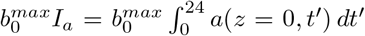, when 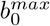 is kept constant. For the sake of comparison, panels (C) and (D) reproduce the results of Figure 3, obtained with the cost function from the main text, 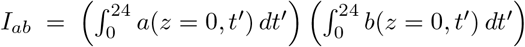. As is apparent, the alternative cost function leads to results that are qualitatively similar to those presented in the main text. Specifically, optimal defense attention can still be either entirely single-channel, e.g., *ν* = 0, or divided to both channels, 0 *≤ ν ≤* 1. Similarly to the results obtained using *I*_*ab*_, when *ϕ*_*a*_ = *ϕ*_*b*_, the optimal defense is *ν* = 0. The ‘spike-protection’ alternative cost function affords the model the same types of optima as with *I*_*ab*_.

Furthermore, one may integrate the cost *I*_*ab*_ over 17 hours instead of 24, parroting the phenomenological period implicit in *ω* = *π/*8.9 hours^*−*1^. In Figure S5 we show heatmaps for *ϕ*_*a*_ versus *ϕ*_*b*_ color-coded with the optimal values of *ν* and *β* which were calculated according to three different cost functions: panels (A-B) repeat the results shown in Figure 4 for comparison, with the cost function being 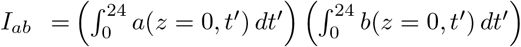 Panels (C-D) follow *I*_*ab*_ but integrate only over the first 17 hours instead of an integration over 24 hours, along the contour 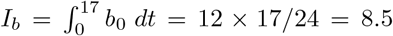 In panels (E-F), we use the ‘spike-protection’ cost function 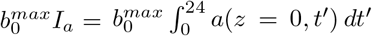, with *ν* and *β* minimizing this cost along the contour 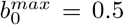, consistent with the average of cos^2^. It is noteworthy that, despite the differences in integration period and bacterial load cost, the resulting optima are qualitatively similar. Therefore, we conclude that the results in this study are not fine-tuned to a specific cost function, but rather, require the minimization of the product of bacterial exposure and AMP levels.

### Appendix F: Reasoning for the AMP boundary conditions

In humans, the mucus is about 200 micrometers thick, but the lumen is measured in centimeters. In mice there are about 20 micrometers of mucus for a lumen width of about 5000 micrometers. Therefore, once the AMP reaches the lumen, which is well mixed because of the watery environment in the small intestine, the AMP concentration is significantly lower than in the mucus. There is strong evidence that AMP secretion does not noticeably affect the bacteria in the lumen [24], therefore, it is reasonable to set *a* = 0 deep in the lumen, and since AMP is not produced in the lumen, it has to be about zero at the boundary, therefore, *a*(*z* = 1) = 0. In contrast, AMP is only produced at *z* = 0 and so it must be removed at *z* = 1, otherwise it would accumulate. Therefore, when there are no other sources of AMP removal, we cannot set *a*^*′*^(*z* = 1) = 0. We are prevented from using this boundary condition. In summary, *a*(*z* = 1) = 0 is the most sensible option.

**FIG. S4.**
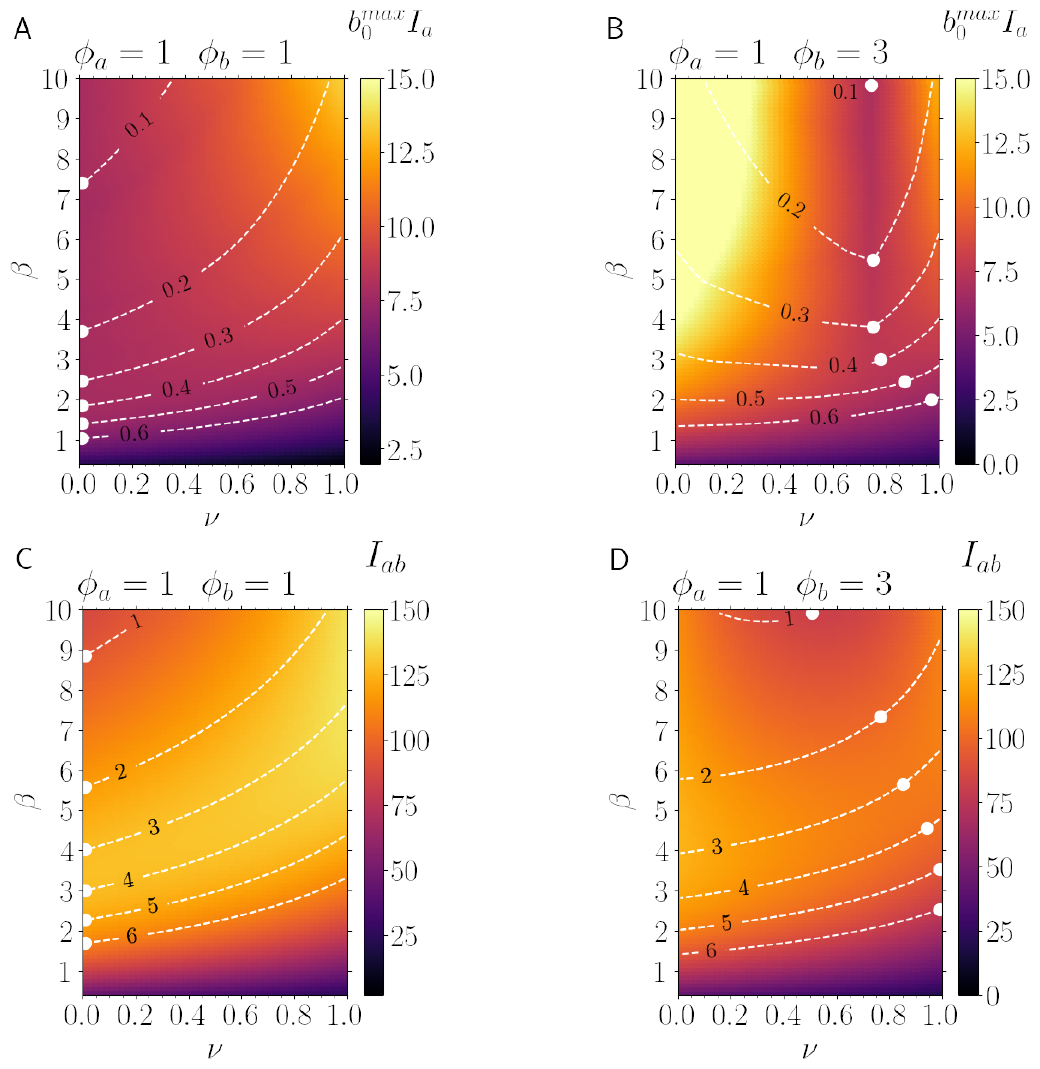
Alternative cost function. Heatmaps for *β* versus *ν* showing optimal host defense. For a fixed maximum epithelial (*z* = 0) concentration of bacteria over a 24h period, 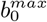, dashed lines indicate equal-contour lines, 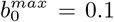, and 0.6, from top to bottom. The minimal host cost in our model is the *β* and *ν* pair on this contour that minimize the cost 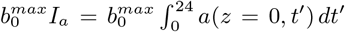, shown as a white dot. (A) *ϕ*_*a*_ = 1, *ϕ*_*b*_ = 1, the SFB anticipate the LPS perfectly. Accordingly, the minimal cost is at *ν* = 0. (B) *ϕ*_*a*_ = 1, *ϕ*_*b*_ = 3, the lumen microbiota lag significantly behind the SFB, and the optimal *ν* involves listening to both channels, 0 *< ν <* 1. (C-D) For comparison, the results in Figure 3, obtained with the cost function *I*_*ab*_. Results for *δt* = 7.2 hours, *ω* = *π/*8.9 hours^*−*1^, *b*_*M* 1_ = 1, *D* = *D*_*A*_ = 1, and v_1_*/μ*_1_ = 1.

**FIG. S5.**
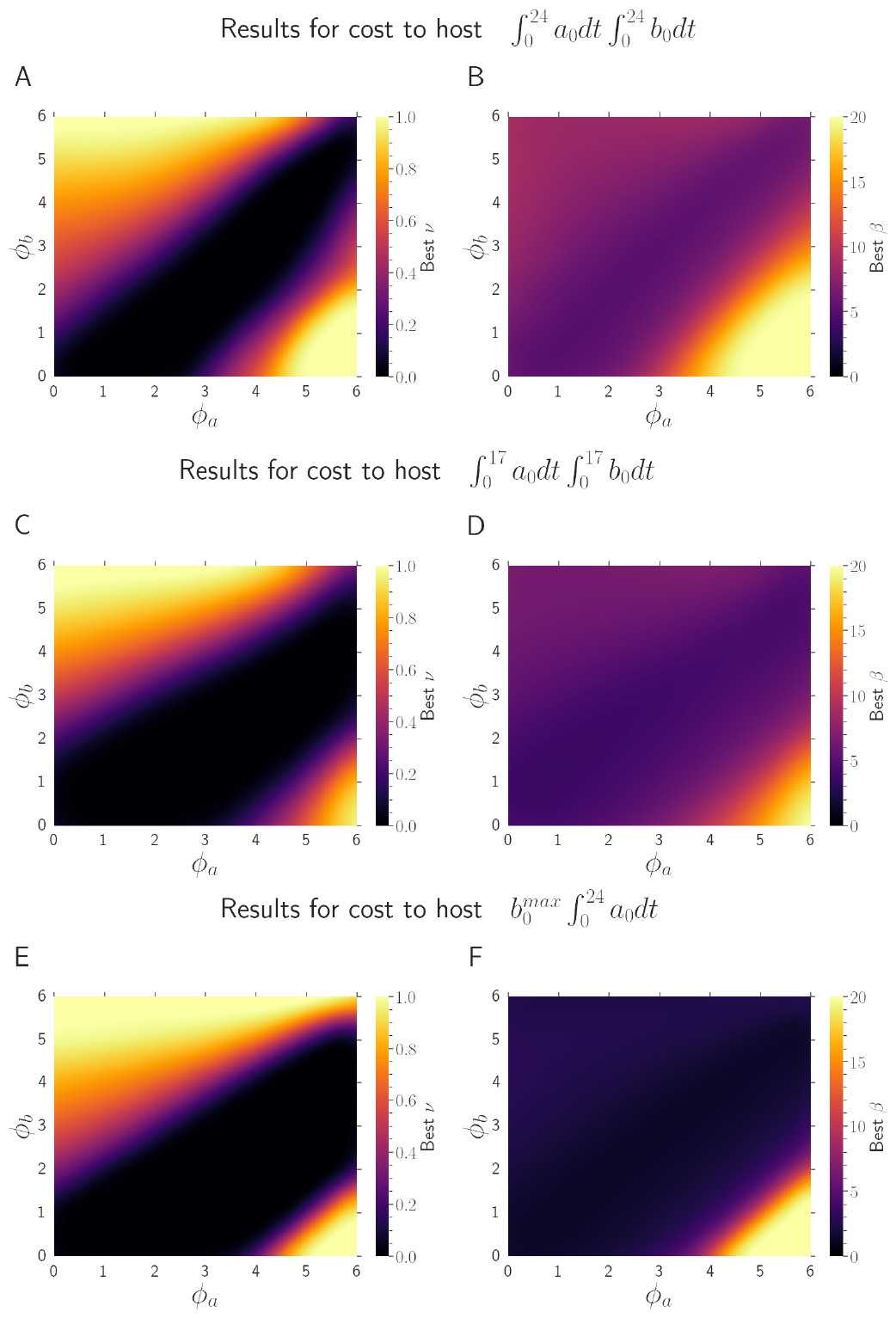
Optimal host response: comparing the effect of different cost functions. (A-B) For comparison, the results from Figure 4, obtained by integrating *I*_*ab*_ for the 24h period. Heat-maps for *ϕ*_*a*_ versus *ϕ*_*b*_ color-coded according to the optimal values of *ν* and *β*. The contour chosen was for *I*_*b*_ = 12. (C-D) Optimal cost when integrating *I*_*ab*_ over 17 hours instead of 24 hours, along the contour 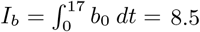 8.5, corresponding to 12 *×* 17*/*24. (E-F) Heat-maps for *ϕ*_*a*_versus *ϕ*_*b*_ color-coded according to the optimal values of *ν* and *β* (E,F respectively); the optimal *ν* and *β* values correspond to the coordinates of the points that minimize the cost function 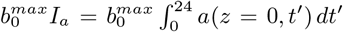 along the contour 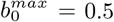, corresponding to the average of *cos*^2^ over its period. All results were obtained for *b*_*M* 1_ = 1, *D* = *D*_*A*_ = 1, and v_1_*/μ*_1_ = 1.

## Notes

https://github.com/AmirErez/GutConjugateDiffusion

### Competing Interest Statement

The authors have declared no competing interest.

### Summary of Updates

Added supplementary figures exploring other cost functions and some explanatory text.

